# Social status in mouse social hierarchies is associated with variation in oxytocin and vasopressin 1a receptor densities

**DOI:** 10.1101/566067

**Authors:** Won Lee, Lisa C Hiura, Eilene Yang, Katherine A Broekman, Alexander G Ophir, James P Curley

## Abstract

The neuropeptides oxytocin and vasopressin and their receptors have established roles in the regulation of mammalian social behavior including parental care, sex, affiliation and pair-bonding, but less is known regarding their relationship to social dominance and subordination within social hierarchies. We have previously demonstrated that male mice can form stable linear dominance hierarchies with individuals occupying one of three classes of social status: alpha, subdominant, subordinate. Alpha males exhibit high levels of aggression and rarely receive aggression. Subdominant males exhibit aggression towards subordinate males but also receive aggression from more dominant individuals. Subordinate males rarely exhibit aggression and receive aggression from more dominant males. Here, we examined whether variation in social status was associated with levels of oxytocin (OTR) and vasopressin 1a (V1aR) receptor binding in socially relevant brain regions. We found that socially dominant males had significantly higher OTR binding in the nucleus accumbens core than subordinate animals. Alpha males also had higher OTR binding in the anterior olfactory nucleus, posterior part of the cortical amygdala and rostral lateral septum compared to more subordinate individuals. Conversely, alpha males had lower V1aR binding in the rostral lateral septum and lateral preoptic area compared to subordinates. These observed relationships have two potential explanations. Preexisting individual differences in the patterns of OTR and V1aR binding may underlie behavioral differences that promote or inhibit the acquisition of social status. More likely, the differential social environments experienced by dominant and subordinate animals may shift receptor expression, potentially facilitating the expression of adaptive social behaviors.

**Highlights:**

- Mice living in social hierarchies express different levels of oxytocin receptor (OTR) and vasopressin 1a receptor (V1aR) binding in various brain regions according to their social status.
- Alphas and subdominants have higher OTR binding in the nucleus accumbens compared to subordinates.
- Alphas have higher OTR binding in the anterior olfactory nucleus compared to subdominants and subordinates.
- Alphas have higher OTR and lower V1aR binding in the rostral lateral septum compared to subordinates.
- Alphas have lower V1aR binding in the lateral preoptic area compared to subordinates.

## Introduction

For socially living species, the ability to modulate behavior across social contexts is essential for survival. Behaving in a socially contextually appropriate manner requires instant decision-making, with individuals having to synchronize information related to their own internal state, their past social experience and their social partner (Chen and Hong, 2018). Social dominance hierarchies are a common form of social organization that requires individuals to adaptively modulate their social behaviors. Hierarchies emerge as unfamiliar individuals encounter and evaluate their opponents’ competitive ability with individuals learning to consistently yield to relatively more dominant social partners (Drews, 1993). Once formed, individuals maintain broadly consistent patterns of both social (e.g. agonistic, submissive, social approach) and non-social (e.g. feeding, drinking, locomotion) behaviors based on their past social interactions (Chase and Seitz, 2011; Fernald and Maruska, 2012; Oliveira and Almada, 1996; Williamson et al., 2016). Animals may however also plastically shift their behavior within a hierarchy should their social status change as a function of a shift in group composition or other ecological or demographic changes (Maruska and Fernald, 2010; Snyder-Mackler et al., 2016; Williamson et al., 2018).

Two neuroendocrine systems that may potentially mediate both stable and adaptive changes in social behaviors within social hierarchies are the neuropeptides oxytocin (OT) and arginine vasopressin (AVP). OT is synthesized in the paraventricular (PVN) and supraoptic nuclei (SON) of the hypothalamus. AVP is also synthesized in the suprachiasmatic nucleus in addition to the PVN and SON. Both hormones are released centrally and are transported to the posterior pituitary where they are released into the periphery. In the brain, OT and AVP are neuropeptides that primarily act on the OT and V1a receptors respectively to regulate social behaviors including maternal behavior (Champagne et al., 2001; Olazábal and Young, 2006), social recognition and memory (Bielsky et al., 2005; Hodges et al., 2017; Veenema et al., 2012), social reward (Dölen et al., 2013; Hung et al., 2017), pair bonding (Ross et al., 2009; reviewed in Young et al., 2011), mating (Ophir et al., 2012, 2008) and aggression (Beiderbeck et al., 2007; Calcagnoli et al., 2014a; Harmon et al., 2002). Individual differences in OTR and V1aR receptor distributions have also been associated with variation in mating strategies and social organization both across (Beery et al., 2008; Insel et al., 1994; Kalamatianos et al., 2010) and within species (Ophir et al., 2012, 2008).

There is some evidence to suggest a relationship between V1aR and OTR signaling and social status. For instance, dominant male rats have been reported to have lower V1aR density in the lateral septum (Askew et al., 2006) while dominant male Syrian hamsters have higher V1aR density in the lateral ventromedial hypothalamus (Cooper et al., 2005). In naked mole rats, dominant queens have lower OTR density in the medial amygdala (MeA) compared to all other animals in each colony (Mooney et al., 2015). The effects of endogenous peptides on behaviors may also be dependent on social status. Dominant, but not subordinate, titi monkeys show increases in sexual and aggressive behavior following central administration with oxytocin (Winslow and Insel, 1991). Likewise, dominant but not subordinate hamsters flank-mark following infusion of oxytocin to the medial preoptic anterior hypothalamic continuum (Harmon et al., 2002). Resident naked mole-rat workers show higher activation of OT neurons in the PVN than soldiers during social encounters (Hathaway et al., 2016) and similar activational differences are observed between dominant and subordinate zebra finches (Kelly and Goodson, 2014) and mandarin voles (Qiao et al., 2014).

Previously, we have demonstrated that outbred CD-1 male mice living in groups of 12 in a large, complex environment rapidly establish linear and stable social dominance hierarchies (So et al., 2015; Williamson et al., 2016). Once hierarchies are established, each mouse holds a unique social rank and shows appropriate agonistic or subordinate behavior to mice of relatively lower or higher ranks (Curley, 2016a; Lee et al., 2018, 2017). We have found that these mice can be subdivided into three broad social status categories. Individuals with the highest social rank are alpha males. These males rarely lose fights and initiate a significant percentage of all agonistic interactions that occur during group housing. Subdominant males receive aggression from more dominant opponents but still actively engage in fights and win consistently over relatively subordinate males. Subordinate individuals are less likely to initiate fights than alpha or subdominant mice and lose far more contests than they win. In addition to changes in aggressive behavior, as individuals become dominant they also produce higher levels of major urinary proteins to scent mark with and also increase their feeding and drinking levels (Lee et al., 2018, 2017; Williamson et al., 2017). In this study, we aimed to investigate whether OTR and V1aR density is associated with social status in mice living in social hierarchies by housing eight cohorts of 12 CD-1 male mice in our large, complex vivaria and then using autoradiography to determine the degree of OTR and V1aR binding in brain regions relevant for social behavior.

## Materials and Methods

### (a) Animals and housing

A total of 96 male outbred Crl:CD1 (ICR) mice aged 7 weeks old were obtained from Charles River Laboratories (Wilmington, MA, USA) and housed in groups of three for 11-15 days in standard sized cages (27 × 17 × 12 cm) with pine shaving bedding. All mice were assigned with unique IDs and marked accordingly by dying their fur with nontoxic animal markers (Stoelting Co., Wood Dale, IL, USA). At 9 weeks of age, mice were weighed then placed into custom-built vivaria as social groups of 12 males and housed for 18-24 days. As previously described (So et al., 2015; Williamson et al., 2016), the vivarium (**Supplemental Figure S1**; 150 × 80cm and 220cm high; Mid-Atlantic, Hagerstown, MD, USA) consists of an upper level with three floor shelves (36,000 cm^2^ = 3 floor × 150 cm × 80 cm) and a lower level with five nest-boxes (2,295 cm^2^ = 5 cages × 27 cm × 17 cm) connected by tubes and all surfaces were covered with pine shavings. Mice can access all levels of the vivarium. We provided standard chow and water ad libitum at the top left and right side of the vivarium. Mice were kept under a 12:12 light:dark cycle with room temperature (22-23°C) and humidity (30-50%) held constant. Within each vivarium, each male had prior social experience with up to only one other male. Trained observers recorded the winner and loser in each instances of agonistic interactions by tallying behaviors such as fighting, chasing, mounting, subordinate posture, and induced-flee behaviors (see **Supplemental Table S1** for ethogram) using handheld Android devices. Data were collected and directly uploaded to Google Drive with timestamps. All observations took place during the dark cycle under red light. We monitored each day of the group housing period if any mouse exhibited a sign of injury or pain. We conducted all procedures with approval from the Columbia University Institutional Animal Care and Use Committee (IACUC protocols: AC-AAAP5405, AC-AAAG0054).

### (b) Tissue collection and preparation

On the day of final observation, mice were euthanized via decapitation at 1400 hours (2h after the light has turned off). Brains were immediately removed then were flash frozen with hexane cooled with dry ice. The brains were stored at −80°C until Cryosectioning. We selected four brains from each cohort (alpha male, 1 subdominant male, and 2 subordinate males) to cut in a cryostat into five sets of 16um coronal sections (80um apart). Sections were mounted on Superfrost Plus slides (Fisher Scientific, Pittsburgh, PA) and stored at −80°C until processing for receptor autoradiography.

### (b) Autoradiography and image analysis

We labeled two of the five sets of slides with ^125^I-labeled radioligands to visualize oxytocin receptor (ornithine vasotocin analogue ([^125^I]-OVTA); NEX254, PerkinElmer, Waltham, MA, USA) and vasopressin 1a receptor (vasopressin (Linear), V-1A antagonist (Phenylacetyl1,0-Me-D-Tyr2,[^125^I-Arg6]-); NEX310, PerkinElmer) as described in Ophir et al. (2013). The radiolabeled slides and ^125^I-labeled radiographic standards (American Radiolabled Chemicals, St Louis, MO, USA) were exposed to phosphor imaging screens (Fujifilm Corporation, Tokyo, Japan) for 2 days. The screens were digitized on a Typhoon FLA 7000 Laser scanner (GE Healthcare, Little Chalfont, UK). Using ImageJ (NIH, Bethesda, MD, USA), we measured optical density in areas of interest and adjacent areas with no receptor binding from three consecutive brain sections bilaterally (except for V1aR in ventral tegmental area) by encircling each region of interest. The optical density values of each structure were averaged then converted to standardized values of disintegration per minute adjusted for tissue equivalent (dpm/mg TE) based on radiographic standards. Specific binding was calculated by subtracting an average of non-specific binding measurements taken from the same brain sections as each region of interest. OTR density was measured between Bregma +2.58 and +1.98 mm in anterior olfactory nucleus (AON), between Bregma +1.78 and +1.34 mm in NAcc (nucleus accumbens core), between Bregma +0.26 and +0.02 mm in LS (lateral septum) and PiR (piriform cortex), and between Bregma −1.58 and −2.06 mm in the posterior ventral medial amygdala nucleus (MeApv), posterior lateral cortical amygdala nucleus (COApl), and basolateral amygdala nucleus (BLA). V1aR density was measured between Bregma +0.98 and +0.74 mm in LSr (rostral lateral septum), medial septal nucleus (MS), vertical limb of the diagonal band (VDB), horizontal limb of the diagonal band (HDB), and ventral pallidum (VP), between Bregma +0.26 and +0.02 mm in LSc (rostral lateral septum) and LPO (lateral preoptic area), and between Bregma −2.92 and −3.16 mm in ventral tegmental area (VTA).Regions were selected based on each of these regions showing high levels of OTR or V1aR binding. Localization of binding in each region was based on the methods of Dubois-Dauphin et al. (1996).

### (c) Statistical analysis

All statistical analyses were undertaken in R version 3.5.1. (Team, 2013).

#### Analysis of agonistic behavior data

The total number of wins and losses from all observed agonistic interactions experienced by each mouse over group housing period was aggregated into frequency win/loss sociomatrices for each cohort. From the sociomatrices, we confirmed the presence of a linear social hierarchy in each cohort by calculating the Landau’s modified *h’* and triangle transitivity (*ttri*) of each cohort and associated p-values derived from 10,000 Monte Carlo randomization (De Vries, 1995; McDonald and Shizuka, 2012) using the compete R package (Curley, 2016b) (see Williamson, Lee & Curley (Williamson et al., 2016) for a more detailed description). Ranks of each mouse in each cohort were determined through calculation of Glicko ratings using the PlayerRatings R package (Stephenson and Sonas, 2012). Briefly, all mice in each cohort are initially assigned with the same rating (2,200 points) then gain or lose points following each agonistic interaction based on the rating difference between themselves and their opponent (Glickman, 1999; Williamson et al., 2016). Based on our behavioral observation of mice social hierarchies, we further categorize individuals in three social status groups: alpha, subdominant and subordinate status groups (Lee et al., 2018, 2017). An alpha male is an individual with social rank 1, with the highest Glicko rating. Mice in the subdominant social status holds Glicko ratings higher than or equal to initially assigned points (2,200) but not the highest rating. The rest of the males in the hierarchy with Glicko ratings lower than 2,200 are categorized as subordinate social status. We calculated the despotism of each alpha male by determining the proportion of all wins by the alpha to all agonistic interactions over the entire group housing period (Williamson et al., 2016). We tested the association between final social rank and body weight measured on the group housing Day 1 for each cohort using Spearman Rank correlation tests and corrected significance values to control for false discovery rate using the Benjamini-Hochberg method (Benjamini and Hochberg, 1995).

#### Analysis of OTR and V1aR density

We analyzed the data on OTR and V1aR density using Bayesian hierarchical linear models fitted in STAN (Carpenter et al., 2017) via the brms R package (Bürkner, 2018). We conducted model selection among combinations of two predictors – social status group (alpha, subdominant, and subordinate) and despotism of each cohort – using leave-one-out cross-validation information criteria (LOOIC) (Gelman et al., 2014). Adding despotism of each cohort as a predictor did not result in a better fit, thus we did not include a predictor of group despotism in the final model for each brain region. To test if mice in different social status groups differ in receptor density, we specified receptor density of each brain region as the outcome variable, social status group as a fixed effect, and imaging screen ID and cohort ID as random effects against a gaussian distribution. With default priors set in the brms package, a model for each brain region and receptor type was fitted with four Markov chains of 2,000 iterations and thinning interval of five. We assessed convergences of the models by checking Gelman-Rubin convergence statistic (Gelman and Rubin, 1992) (all models at convergence, Rhat=1) and by visually inspecting MCMC objects. Statistical significance of differences in receptor density among three social status groups can be inferred when 95% credibility intervals (CI) do not overlap zero.

## Results

### (a) Social hierarchy characteristics

All 8 cohorts of 12 males formed significantly linear dominance hierarchies (h’ mean = 0.80, interquartile ranger (IQR) = [0.74 – 0.89], all p<0.01) with significantly high triangle transitivity (*ttri* = 0.88 [0.85 – 0.94], all p<0.001) and directional consistency of aggressive behavior (DC = 0.88 [0.85 – 0.90]). The mean degree of alpha male despotism over the entire group housing period was 0.47 [0.38 – 0.52]. Body weight measured on Day 1 of the group housing period was not significantly associated with final social rank in any social group (Spearman’s rank correlation tests, controlled for FDR, all p>0.68).

### (b) Oxytocin receptor (OTR) density

OTR density differences across social status groups were found in 4 of 7 regions analyzed (see Fig. 1a for representative images, Fig. 1b for boxplots with raw data points and Fig. 1c for statistical comparisons for each brain region). In the AON, OTR density was significantly higher in alpha males than in both subdominant and subordinate males (b_subdominant-alpha_: −486 dpm/mg TE [−905, −113]; b_subordinate-alpha_: −627 dpm/mg TE [−985, −282]) but there was no OTR density difference between subdominant and subordinate males. In the NAcc, both alpha and subdominant mice had higher OTR than did subordinate (b_subordinate-alpha_: −980 dpm/mg TE [−1430, −507]; b_subordinate-subdominant_: −930 dpm/mg TE [−1342, −459]) while alpha and subdominant males did not differ in OTR density. In the LS and the COApl, alpha males showed higher OTR density than did subordinate males (LS – b_subordinate-alpha_: −301 dpm/mg TE [−535, −82]; COApl – b_subordinate-alpha_: −391 dpm/mg TE [−720, −97]). There was no notable difference among alpha, subdominant and subordinate males in the PiC, MeA, and BLA.

**Figure 1.**
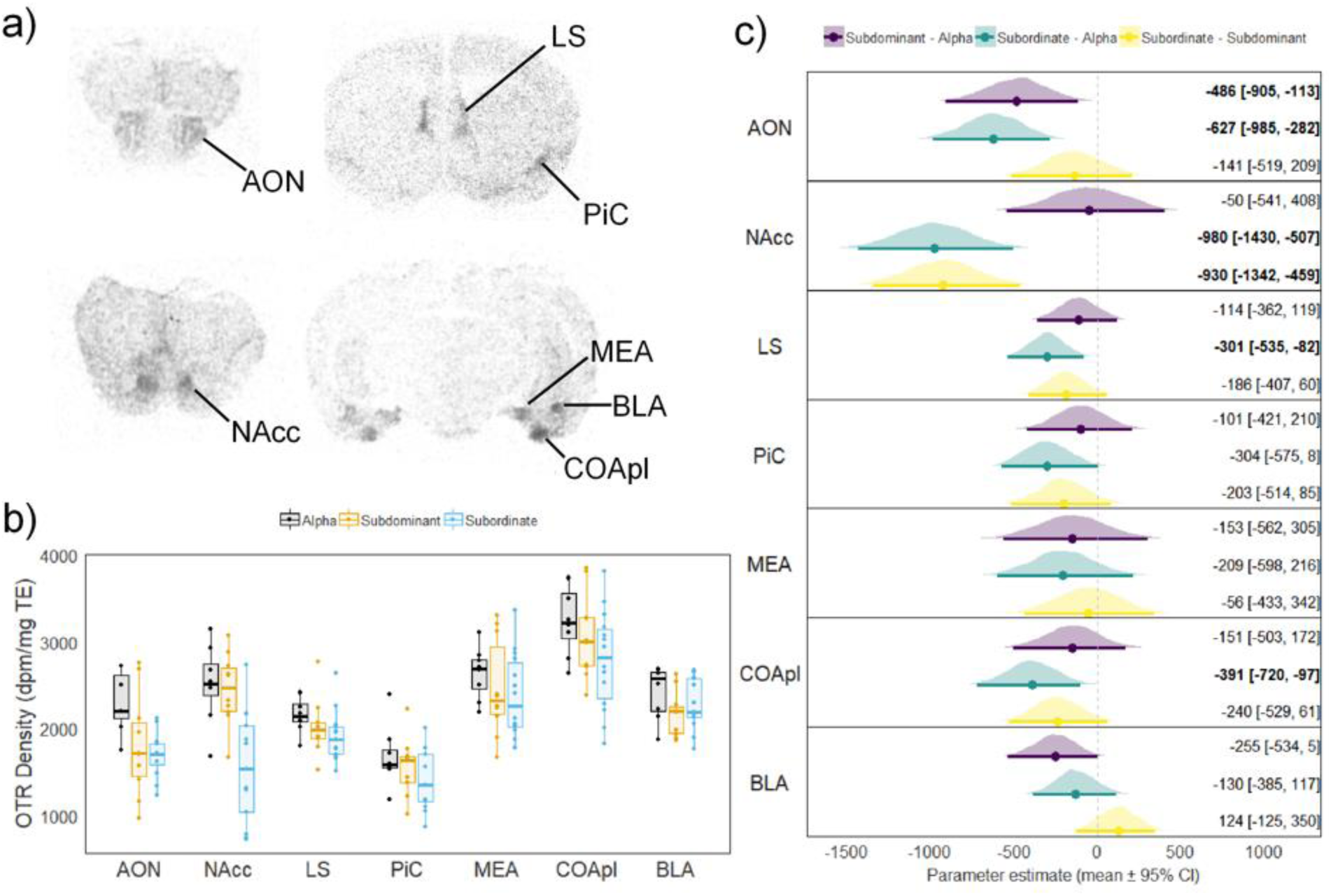
OTR density by social status groups in seven brain regions analyzed. (a) Representative autoradiographs of OTR binding in coronal sections. Measured brain regions are labeled. (b) Raw data and boxplots of OTR density by social status groups in each brain region. Boxplots show median (horizontal bars), interquartile ranges (boxes) and 95% confidence intervals (whiskers). (c) Posterior densities, posterior means (solid dots) and 95% credible intervals (bars) for comparison among social status groups in each brain region. Text on the right notes mean estimates and 95% CI for each comparison.

### (c) Vasopressin subtype 1a receptor (V1aR) density

V1aR density differences across social status groups were found in 2 of 7 regions analyzed (see Fig. 2a for representative images, Fig. 2b for boxplots with raw data points and Fig. 2c for statistical comparisons for each brain region). Alpha males had lower V1aR density than did subordinate males in the LSr (b_subordinate-alpha_: 762 [202, 1358]) and in the LPO (b_subordinate-alpha_: 666 [40, 1291] dpm/mg TE). There was no notable difference among alpha, subdominant and subordinate males in any of the other brain regions analyzed.

**Figure 2.**
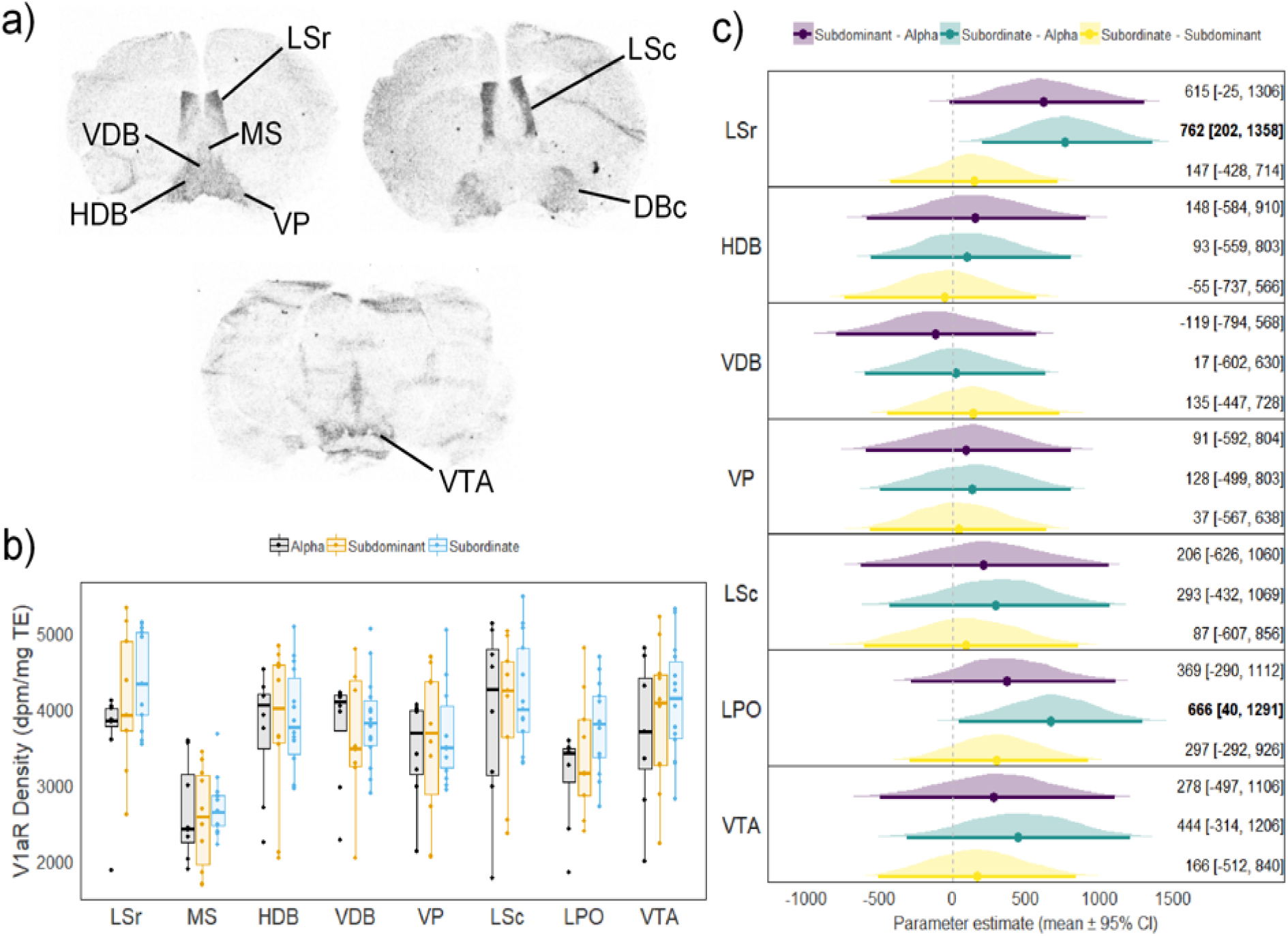
V1aR density by social status groups in seven brain regions analyzed. (a) Representative autoradiographs of V1aR binding in coronal sections. Measured brain regions are labeled. (b) Raw data and boxplots of V1aR density by social status groups in each brain region. Boxplots show median (horizontal bars), interquartile ranges (boxes) and 95% confidence intervals (whiskers). (c) Posterior densities, posterior means (solid dots) and 95% credible intervals (bars) for comparison among social status groups in each brain region. Text on the right notes mean estimates and 95% CI for each comparison.

## Discussion

In the current study we demonstrate that the social status of mice living in social hierarchies is associated with individual variation in oxytocin and vasopressin 1a receptor binding density. We were able to characterize three categories of male mice: i) alpha males that show consistently higher levels of aggression and territorial behavior and rarely lose any contests; ii) subdominant males that show high levels of aggression towards less dominant individuals but that also receive aggression from the alpha male and other more dominant males; iii) subordinate males that rarely win any fights and consistently lose to many other more dominant males. We found that alpha males had significantly higher oxytocin receptor densities than subordinate males in the LS and COApl. Alpha males also had significantly higher oxytocin receptor densities than both subdominants and subordinates in the AON. The largest effect size was observed in the NAcc, where both alpha and subdominant individuals had higher OTR binding than subordinates. No differences between social statuses in OTR binding were observed in the PiC, MeA or BLA. Fewer group differences were observed in vasopressin 1a receptor densities with alpha males found to have significantly lower V1aR densities than subordinate males only in the LPO and LSr.

The observed increases in OTR in several brain regions of more dominant males are notable as generally an up-regulation of oxytocinergic signaling is associated with reduced aggression in male rodents. Rats selectively bred for high levels of inter-male aggression have lower levels of OT mRNA in the PVN (Calcagnoli et al., 2014a), less aggressive male mice have more OT neurons in the same region (Tsuda et al., 2011), and non-aggressive worker naked mole-rats show increased activation of PVN OT neurons in response to intruders whereas aggressive soldiers do not (Hathaway et al., 2016). Microdialysis studies show that OT is released from male rats during resident-intruder aggression (de Jong and Neumann, 2018). Variation in oxytocin receptors has also been associated with aggressive phenotypes in rats. WTG rats show higher levels of OTR binding in the central nucleus of the amygdala (CeA) and the bed nucleus of the stria terminalis (BNST) but not the LS compared to less aggressive strains. Reduced OTR binding in the caudate putamen and LS but elevated OTR binding in the medial preoptic area (mPOA) have been reported in aggressive compared to non-aggressive Wistar rats (Lukas et al., 2010). Transgenic mice that do not possess any oxytocin receptors exhibit significantly higher aggression than wild-type animals (Dhakar et al., 2012; Hattori et al., 2015; Sala et al., 2013, 2011; Takayanagi et al., 2005), unless the ablation is limited to serotonergic neurons in the dorsal and median raphe nucleus which leads to reduced aggression (Pagani et al., 2015). Moreover, in both rats and mice central or intransal infusion of OT ameliorates aggression while infusion of OTR antagonists elevates male aggression (Arakawa et al., 2015; Calcagnoli et al., 2013, 2014b, 2015a; Karpova et al., 2016). Region specific infusion of OT into the CeA of rats (Calcagnoli et al., 2015b) and mPOA and anterior hypothalamus (AH) of Syrian hamsters (Harmon et al., 2002) also reduced aggression.

There are several possible reasons why we observe elevated oxytocin receptor binding in the brain of more dominant animals. First, although higher OTR densities suggest increased OT neurotransmission it is possible that locally increased OTR binding may be indicative of reduced OT neurotransmission due to reduced ligand availability (de Jong and Neumann, 2018). Second, most of the studies establishing a relationship between reduced oxytocinergic signaling and higher aggression investigate aggression in short dyadic resident-intruder paradigms. Such aggression is not necessarily equivalent to the dominance behavior that is established over several days in a group living context. It is possible that the oxytocinergic system is used in a different manner to regulate social relationships over longer time periods. Consistent with this hypothesis, in a study that examined the establishment of social hierarchies, subordinate, but not dominant, rats were found to have reduced OTR mRNA expression in the MeA following aggressive and stressful interactions (Timmer et al., 2011). Thirdly, the observed receptor binding differences may reflect experience-dependent changes in the expression of receptors induced via the differential social environments experienced by dominant and subordinate animals. Previous work has established that variation in social environments can induce such changes in the distribution of both OTR and V1aR (Curley et al., 2009; Hiura and Ophir, 2018; Prounis et al., 2015; Winslow et al., 1993). In addition to changes in aggressive and subordinate behaviors displayed by mice of different social ranks, individuals also show plastic changes in several other behaviors. We have previously shown that during the acquisition of social ranks, dominant males increase their patrolling, feeding, drinking and urination behaviors (Lee et al., 2018, 2017; Williamson et al., 2016) whereas subordinate animals become less active and avoid the dominant males (Curley, 2016a; Lee et al., 2018). In mice, differences in OTR binding may reflect plastic changes in other behavioral and physiological phenotypes related to social status.

The biggest effect observed in this study was the increased OTR binding density in the NAcc of both alpha and subdominant animals compared to subordinates. Classically, a role for oxytocin in regulating pair-bonding and parental behavior has been established in prairie voles. Monogamous prairie voles have higher levels of OTR in the NAcc compared to promiscuous meadow voles (Insel and Shapiro, 1992), oxytocin receptor density in the OTR is associated with spontaneous parental care in female prairie voles (Olazábal and Young, 2006) and sexual fidelity in male prairie voles (Ophir et al., 2012) and the action of oxytocin on these receptors facilitates pair-bonding (Liu and Wang, 2003; Ross et al., 2009) and promotes social novelty seeking behavior in adolescent male rats (Smith et al., 2017). In mice, there does exist a direct axonal oxytocinergic projection from the PVN to the NAcc (Dölen et al., 2013). It has recently been shown that NAcc oxytocin receptors in male mice are required for conditioned place preference to social stimuli but not to cocaine, suggesting that oxytocin in this region regulates social reward (Dölen et al., 2013). Notably, aggressive CD-1 male mice show conditioned place preference for contexts in which they have experienced being aggressive which lasts for several days (Golden et al., 2017). These mice will also work via lever pressing or nose pokes to attack subordinate males under several reinforcement schedules (Couppis and Kennedy, 2008; Falkner et al., 2016; Golden et al., 2017) as well as relapsing to aggression seeking following forced or voluntary abstinence (Golden et al., 2017). It is possible therefore that the increased OTR binding in the NAcc of dominant animals in our study is related to an increased motivation to access and dominate more subordinate animals, which could function to monitor their social environment (territory) for competitors and enforce their dominant status. Congruent with this explanation, male mice that are socially defeated show consistent social withdrawal behavior (Berton et al., 2006). Chronically defeated mandarin voles also show reduced levels of OT and OTR in the NAcc and their social withdrawal can be reversed with microinjection of OT into this region (Wang et al., 2018). In social hierarchies, it is possible that the reduced levels of OTR binding facilitate reduced social interaction, which may be beneficial for subordinate males to reduce the number of potentially harmful interactions with more dominant individuals.

We also observed higher OTR binding in alpha males compared to all other males in the anterior olfactory nucleus and compared to subordinate males only in the COApl. Both regions, along with the PiC, where we did not observe any differences between social ranks in oxytocin receptor binding, receive projections from the main olfactory bulb, which primarily processes volatile odors (Wacker and Ludwig, 2012). Conversely, the MeA, which did not differ in oxytocin receptor binding between ranks, receives relatively few projections from the main olfactory bulb but many from the accessory olfactory bulb. The AON, which receives dense OT projections from the PVN (Knobloch et al., 2012), has a well-established role in the processing of social and olfactory information, including oxytocin receptor dependent social recognition (Wacker and Ludwig, 2012). Optogenetically evoked oxytocin release in the AON increases the olfactory investigation of same-sex conspecifics and enhances social recognition, whereas the inhibition of oxytocin receptors in this region inhibits same-sex social recognition (Oettl et al., 2016). Indeed, it has been argued that oxytocin acting on its receptors in this region are part of a social-salience brain network whereby the salience of social stimuli are increased such that relevant social information is extracted and integrated with other information during social encounters (Johnson et al., 2017). Recent work has shown that in addition to its role in regulating social recognition, the COA also possesses distinct cell populations that assess and respond to the saliency of both appetitive and aversive social odors (Root et al., 2014) and is activated during social interaction (Kim et al., 2015). In particular, activation of the posterior part of the COA stimulates response towards attractive volatile odors include those of other males (Root et al., 2014). This region also possesses high numbers of estrogen receptors suggesting that changes in oxytocin receptor density might be initiated by hormonal changes following differential social experiences (Cai et al., 2014). Taken together, it seems possible that the increased levels of oxytocin receptors in the AON and COApl may lead to increased social interaction by dominant males by enhancing the saliency of volatile odors from other males.

In the rostral LS, we observed that dominant alpha males had significantly higher levels of OT binding and lower levels of V1aR binding compared to subordinate males. There was no difference in V1aR binding in the caudal LS between dominants and subordinates. AVP fibers innervating the LS originate from the BNST and have been considered to promote inter-male aggression, although the relationship between septal vasopressin and aggression appears to be complex (Veenema et al., 2010). Aggressive male rats and mice have lower AVP fiber densities in the lateral septum than non-aggressive animals (Beiderbeck et al., 2007; Compaan et al., 1993; Everts et al., 1997), but release of AVP in the lateral septum has been associated with both increased (Veenema et al., 2010) and decreased (Beiderbeck et al., 2007) levels of inter-male aggression. Further, septal AVP infusions typically increase inter-male aggression (Veenema et al., 2010) but may not depending upon context (Beiderbeck et al., 2007). Our findings are congruent with one other study that found that subordinate male rats had higher V1aR binding in the LS than dominant male rats when housed in pairs (Askew et al., 2006). The differential expression of V1aR within the sub-units of the LS that we report might provide some insight into the conflicting results regarding how the LS impacts aggression. The LS projects to the ventromedial hypothalamus, in which sub-units and specific cell types impact aggression differentially (Hashikawa et al., 2016). Furthermore, the LS modulates aggression via VMH projections (Falkner et al., 2016). Therefore, it is possible that different portions of the LS (e.g. rostral or caudal) and different vasopressin-V1aR modulation therein impact agonistic behavior in different ways. Furthermore, it is unclear whether the observed differences in septal V1aR are related to differences in aggressive behavior between dominant and subordinate males, or whether they influence other behavioral processes that are known to be regulated by septal AVP such as anxiety-related behavior (Veenema et al., 2010) and social recognition (Bielsky et al., 2005; Dantzer et al., 1988; Engelmann and Landgraf, 1994; Everts et al., 1997; Landgraf et al., 2003).

Similar to vasopressin, septal oxytocin also modulates social behaviors and its relationship to aggressive and subordinate behaviors are unclear. Contrary to our findings, one study found no differences between high and low aggressive rats in LS OTR binding (Calcagnoli et al., 2014a) and another study found that chronically socially defeated mice actually have elevated OTR mRNA compared to control mice (Litvin et al., 2011). However, increased activation of septal oxytocin receptors is known to promote social recognition and memory (Lukas et al., 2013) and attenuate social fear of conspecifics (Zoicas et al., 2014). It is possible that the observed increases in oxytocin receptor binding of alpha males in these social hierarchies do not relate to changes in agonistic behavior but do relate to the attenuation of social fear compared to subordinate males. Moreover, it is interesting to note that OTR LS and V1aR LSr showed the opposite pattern of binding across hierarchy position (i.e. relatively high OTR and low V1aR in alphas compared to subordinates). A similar relationship between high OTR and low V1aR LS density has been described in relation to social investigation (Ophir et al., 2009). This opposite pattern of receptor density within the LS might further explain the complicated relationship between nonapeptides and the modulation of aggression.

Finally, we also observed higher V1aR binding in the LPO of subordinate males compared to dominant alpha males. The functional significance of this finding is unclear. In hamsters, it is well established that infusion of AVP into an area extending from the mPOA to the anterior hypothalamus including the LPO promotes flank marking, flank gland grooming and inter-male aggression (Albers et al., 1988; Ferris et al., 1984). In the preoptic area of mice, V1aR binding is limited only to the LPO and is not observed in the mPOA (Dubois-Dauphin et al., 1996). Although the functional role of AVP in the LPO in regulating mouse aggression is unclear, AVP infusion into the preoptic area of mice increases self-grooming and decreases locomotion and olfactory investigation of novel stimuli (Lumley et al., 2001). Potentially, the increased V1aR binding observed in subordinate mice may promote their reduced locomotor and investigatory activities compared to dominant males.

## Conclusions

We found that male mice living in large social groups formed highly linear social hierarchies and that individual social status within these hierarchies was associated with differences in the binding levels of oxytocin receptors and, to a lesser extent, vasopressin 1a receptors. There are likely two possible explanations for these observed associations. First, it is possible that preexisting individual differences in the expression of OTR and V1aR in the brain regions we identified between alpha, subdominant and subordinate males may facilitate the ascension or inhibition of social rank. However, given that we observed variation in several brain regions, several of which are not classically related to the promotion of aggressive behavior, this seems unlikely. A more parsimonious interpretation is that as animals acquire their social status, plastic changes in the expression of these receptors, particularly OTR, emerge. These changes may underlie adaptive changes in other social behaviors that occur as animals become dominant or subordinate. For instance, we suggest that increases in OTR binding in the NAcc of dominant animals may facilitate increased social approach behavior to conspecifics. Increased OTR binding in the AON and COApl of dominants may lead to improved olfactory processing of volatile odors whereas increased OTR binding in the rostral LS may inhibit social fear. Future mechanistic experiments will be required to test these hypotheses.

## Acknowledgements

We thank Dr. Frances Champagne for advice and suggestions in writing the manuscript, Jaxon Paige Bowman for assistance in image analysis and Curley Lab students for help with behavioral observations.

## Data availability

All raw data and code used in this paper are publicly available at GitHub: https://github.com/jalapic/otr_v1ar

## Contributions

WL, AGO & JPC conceived and planned the experiments. WL, LCH, EY & KAB carried out the experiments. WL & JPC analyzed the data. WL and JPC took the lead in writing the manuscript. All authors provided critical feedback and helped shape the research, analysis and manuscript.

