## Supplemental tables and figures for "Social status in mouse social hierarchies is associated with variation in oxytocin and vasopressin 1a receptor densities"

**Supplemental Table S1. Mouse social behavior ethogram**

| Priority | Behavior | Description |
|---|---|---|
| 1 | Fighting | Individual lunges at and/or bites the other individual |
| 2 | Chasing | Individual follows the target individual rapidly and aggressively while the other individual attempts to flee |
| 3 | Mounting | Individual mounts another individual from behind with the recipient attempting to flee or otherwise being pinned to the floor |
| 4 | Subordinate posture | Individual responds to the approach from another individual by remaining motionless and/or exposing their nape |
| 5 | Induced-flee | Individual flees without any aggression shown by another individual |

**Supplementary Figures**

**Supplemental Figure S1.** Vivarium for social group housing.

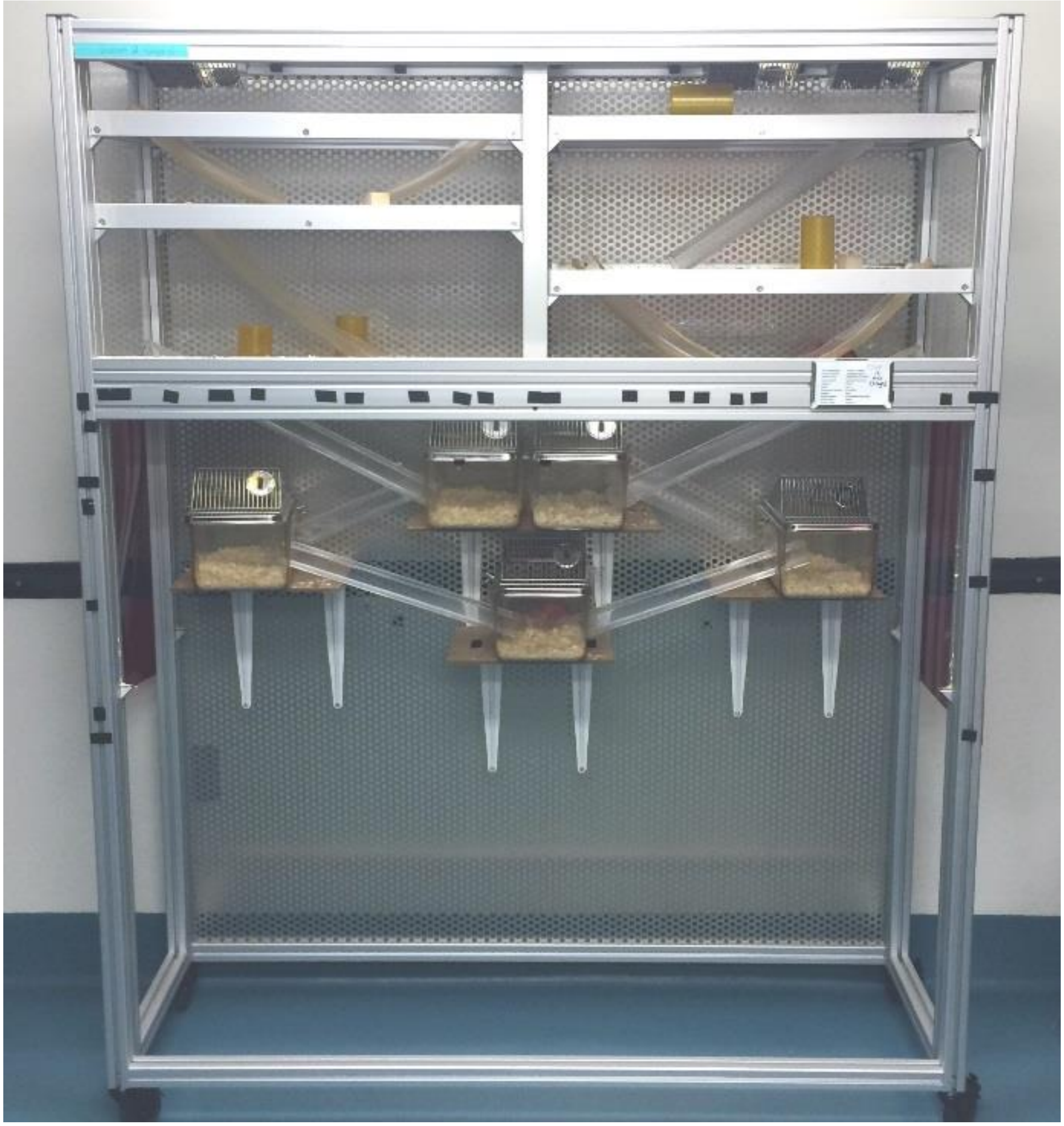
